# Mutational signatures of DNA mismatch repair deficiency in *C. elegans* and human cancers

**DOI:** 10.1101/149153

**Authors:** B Meier, NV Volkova, Y Hong, P Schofield, PJ Campbell, M Gerstung, A Gartner

**Affiliations:** Centre for Gene Regulation and Expression, University of Dundee, Dundee, UK.; European Molecular Biology Laboratory, European Bioinformatics Institute (EMBL-EBI), Hinxton, Cambridgeshire, UK.; Division of Computational Biology, University of Dundee, Dundee, UK.; Cancer Genome Project,Wellcome Trust Sanger Institute, Hinxton, Cambridge, UK.; Department of Haematology, University of Cambridge, Cambridge, UK.; Department of Haematology, Addenbrooke’s Hospital, Cambridge, UK.

**Keywords:** DNA mismatch repair, mutational signatures, hypermutation, replication errors, POLE-4, genome integrity, *C. elegans*

## Abstract

Throughout their lifetime cells are subject to extrinsic and intrinsic mutational processes leaving behind characteristic signatures in the genome. DNA mismatch repair (MMR) deficiency leads to hypermutation and is found in different cancer types. While it is possible to associate mutational signatures extracted from human cancers with possible mutational processes the exact causation is often unknown. Here we use *C. elegans* genome sequencing of *pms-2* and *mlh-1* knockouts to reveal the mutational patterns linked to *C. elegans* MMR deficiency and their dependency on endogenous replication errors and errors caused by deletion of the polymerase ε subunit *pole-4*. Signature extraction from 215 human colorectal and 289 gastric adenocarcinomas revealed three MMR-associated signatures, one of which closely resembles the *C. elegans* MMR spectrum and strongly discriminates microsatellite stable and unstable tumors (AUC=98%). A characteristic difference between human and *C. elegans* MMR deficiency is the lack of elevated levels of N<u>C</u>G>NTG mutations in *C. elegans*, likely caused by the absence of cytosine (CpG) methylation in worms. The other two human MMR signatures may reflect the interaction between MMR deficiency and other mutagenic processes, but their exact cause remains unknown. In summary, combining information from genetically defined models and cancer samples allows for better aligning mutational signatures to causal mutagenic processes.

## INTRODUCTION

Cancer is a genetic disease associated with the accumulation of mutations. A major challenge is to understand mutagenic processes acting in cancer cells. Accurate DNA replication and the repair of DNA damage are important for genome maintenance. The identification of cancer predisposition syndromes caused by defects in DNA repair genes was important to link the etiology of cancer to increased mutagenesis. One of the first DNA repair pathways associated with cancer predisposition was DNA mismatch repair (MMR). MMR corrects mistakes that arise during DNA replication. Mutations in MMR genes are associated with hereditary non-polyposis colorectal cancer (HNPCC), also referred to as Lynch Syndrome (Fishel et al. 1993; Bronner et al. 1994; Nicolaides et al. 1994; Papadopoulos et al. 1994; Miyaki et al. 1997).

DNA mismatch repair is initiated by the recognition of replication errors by MutS proteins, initially defined in bacteria. In *S. cerevisiae* and mammalian cells, two MutS complexes termed MutSα and MutSβ comprised of MSH2/MSH6 and MSH2/MSH3 heterodimers, respectively, are required for DNA damage recognition albeit with differing substrate specificity (Drummond et al. 1995; Habraken et al. 1996; Genschel et al. 1998). Binding of MutS to the DNA lesion facilitates subsequent recruitment of the MutL complex. MutL enhances mismatch recognition and promotes a conformational change in MutS through ATP hydrolysis to allow for the sliding of the MutL/MutS complex away from mismatched DNA (Allen et al. 1997; Gradia et al. 1999). DNA repair is initiated in most systems by a single-stranded nick generated by MutL (MutH in *E. coli*) on the nascent DNA strand at some distance to the lesion (Kadyrov et al. 2006; Kadyrov et al. 2007). Exonucleolytic activities in part conferred by Exo1 contribute to the removal of the DNA stretch containing the mismatch followed by gap filling via lagging strand DNA synthesis (Goellner et al. 2015). The most prominent MutL activity in human cells is provided by the MutLα heterodimer MLH1/PMS2 (Prolla et al. 1998; Cannavo et al. 2005). Moreover, human MLH1 is found in heterodimers with PMS1 and MLH3, called MutLβ and MutLγ. Of these only MutLγ is thought to have a minor role in MMR (Cannavo et al. 2005). The *C. elegans* genome does not encode obvious MutLβ and γ subunits (PMS1 and MLH3 homologues, respectively), while the MutLα subunits MLH-1 and PMS-2 can be readily identified using homology searches (Supplemental Table S1).

Analysis of mutations in microsatellite loci of *MLH1*-deficient colorectal cancer cell lines suggested rates of repeat expansion or contraction between 8.4x10^−3^ to 3.8x 10^−2^ per locus and generation (Bhattacharyya et al. 1994; Hanford et al. 1998). Estimates using *S. cerevisiae* revealed a 100-to 700-fold increase in DNA repeat tract instability in *pms2*, *mlh1* and *msh2* mutants (Strand et al. 1993) and a ~5-fold increase in base substitution rates (Yang et al. 1999). *C. elegans* assays using reporter systems or selected, PCR-amplified regions revealed a more than 30-fold increased frequency of single base substitutions in *msh-6*, a 500-fold increase in mutations in A/T homopolymer runs and a 100-fold increase in mutations in dinucleotide repeats (Degtyareva et al. 2002; Tijsterman et al. 2002; Denver et al. 2005), akin to the frequencies observed in yeast and mammalian cells (Strand et al. 1993; Hanford et al. 1998). Recently, whole genome sequencing approaches using diploid *S. cerevisiae* started to provide a genome-wide view of MMR deficiency. *S. cerevisiae* lines carrying an *msh2* deletion alone or in conjunction with point mutations in one of the three replicative polymerases, Polα/primase, Polδ, and Polε, were propagated over multiple generations to determine the individual contribution of replicative polymerases and MMR to replication fidelity (Lang et al. 2013; Lujan et al. 2014; Lujan et al. 2015). These analyses estimated an average base substitution rate of 1.6 x 10^−8^ per base pair per generation in *msh2* mutants and a further increased rate in double mutants of *msh2* and any of the replicative polymerases (Lujan et al. 2014; Lujan et al. 2015). A synergistic increase in mutagenesis was also recently observed in childhood tumors in which MMR deficiency and mutations in replicative polymerase ε and δ, required for leading and lagging strand DNA synthesis respectively, occurred (Shlien et al. 2015).

In human cancer samples 30 mutational signatures (referred to as COSMIC signatures from here on) have been uncovered by mathematical modeling across a large number of cancer genomes representing more than 30 tumor types (http://cancer.sanger.ac.uk/cosmic/signatures) (Alexandrov et al. 2013a; Alexandrov et al. 2013b). These signatures are largely defined by the relative frequency of the six possible base substitutions (C>A, C>G, C>T, T>A, T>C, T>G) in the sequence context of their adjacent 5’ and 3’ base. Of these, COSMIC signatures 6, 15, 20, 21 and 26, have been associated with MMR deficiency with several MMR signatures being present in the same tumor sample (Alexandrov et al. 2013a; Alexandrov et al. 2013b). It is not clear if these MMR signatures are conserved across evolution and how they reflect MMR defects. Therefore, MMR signatures deduced from defined monogenic MMR defective backgrounds (which we will herein refer to as mutational patterns) could contribute to the refinement of computationally derived mutational signatures extracted from cancer genomes.

Here we investigate the genome-wide mutational impact of the loss of the MutL mismatch repair genes *mlh-1* and *pms-2* in the nematode *C. elegans*. Furthermore, we address the contribution of a deletion of *pole-4*, a non-essential accessory subunit of the leading-strand DNA polymerase Polε, to mutation profiles and hypermutation.

## RESULTS

### Mutation rates and profiles of *mlh-1*, *pms-2* and *pole-4* single mutants grown over 20 generations

We previously established *C. elegans* mutation accumulation assays and demonstrated that defects in major DNA damage response and DNA repair pathways, including nucleotide excision repair, base excision repair, DNA crosslink repair, DNA end-joining and apoptosis did not lead to overtly increased mutation rates when lines were propagated for 20 generations (Meier et al. 2014). The experimental setup takes advantage of the 3-4 days life cycle of *C. elegans* and its hermaphroditic reproduction by self-fertilization. This allows for the propagation of clonal *C. elegans* lines, which in each generation pass through a single cell bottleneck provided by the zygote. We now extend these studies to MMR deficiency conferred by MutLα mutations *mlh-1* and *pms-2*. Given that null alleles of the human and *C. elegans* leading strand polymerase Polε catalytic subunit, *POLE* and *pole-1*, respectively, cause lethality, we focused our analysis on a non-essential *C. elegans* Polε subunit, termed POLE-4. Dbp3p, the *S. cerevisiae* POLE-4 ortholog, has been implicated in stabilizing POLE interaction with the primer-template DNA complex (Aksenova et al. 2010).

We detected an average of 4 base substitutions and 2 insertions or deletions in wild-type *C. elegans* lines propagated for 20 generations (Fig. 1A, 1B). In contrast, *mlh-1* and *pms-2* mismatch repair single mutants carried an average of 1174 and 1191 unique mutations, respectively, of which 288 and 309 were base substitutions (Fig. 1A) and 886 and 882 indels, defined as small insertions and deletions of less than 400 base pairs (Fig. 1B). The nature of single nucleotide changes and the overall mutation burden were congruent across independent lines of the same genotype and mutation numbers linearly increased from F10 to F20 generation lines (Fig. 1). In contrast to *mlh-1* and *pms-2*, *pole-4* mutants exhibited mutation numbers and profiles not significantly different from wild-type (Fig. 1, Supplemental Table S2).

**Figure 1.**
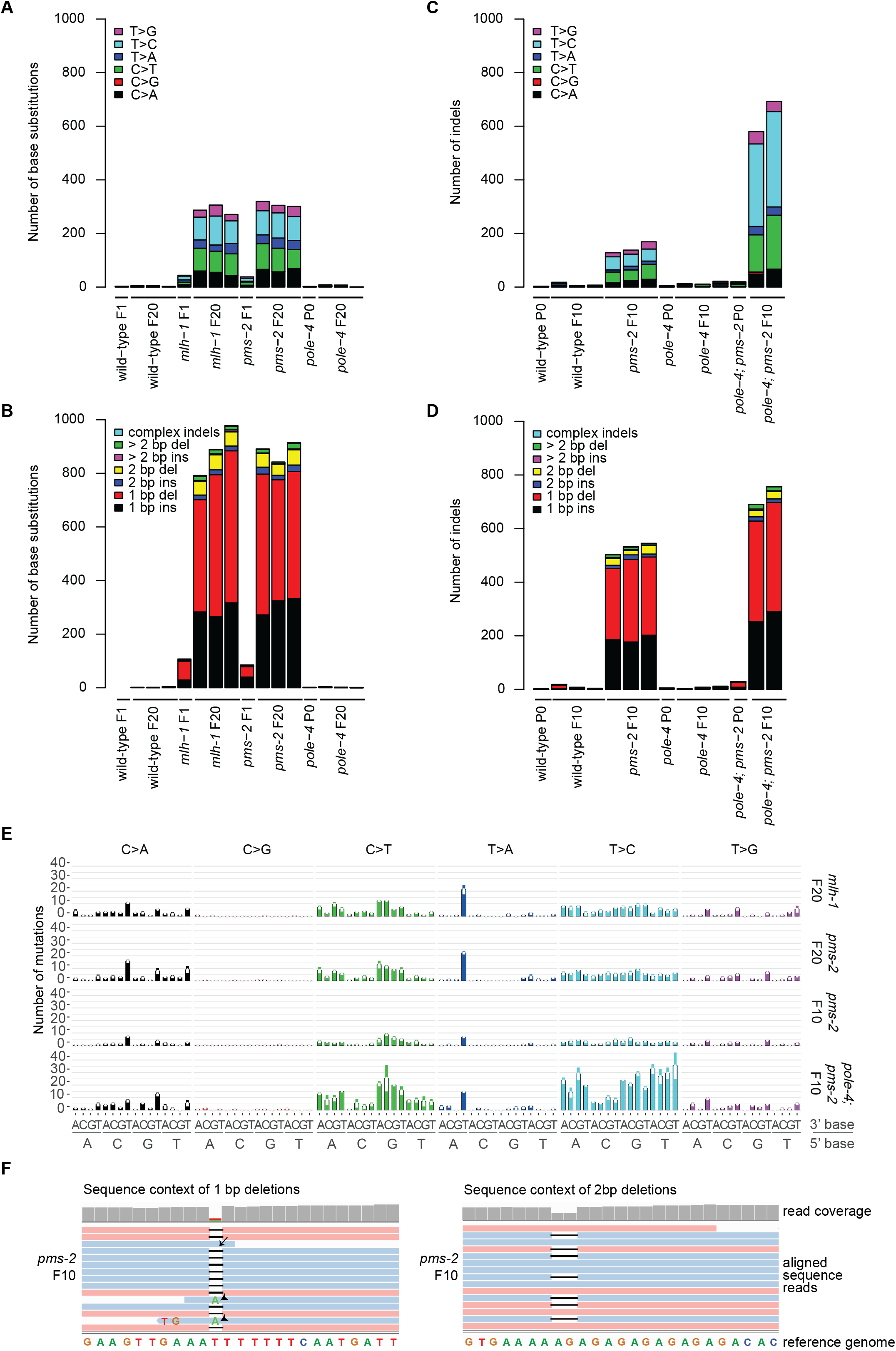
Mutations in *C. elegans* wild-type and mutant strains grown for 10 or 20 generations. Identical base substitutions as well as indels occurring in the same genomic location among samples of the entire dataset (duplicates) were excluded from the analysis, thus only reporting mutations unique to each individual sample. **(A)** Number and types of base substitutions identified in the parental (P0) or one first generation (F1) line and three independently propagated F20 lines of wild-type, *mlh-1, pms-2*, and *pole-4* single mutants. **(B)** Number and types of insertions and deletions (indels) identified in initial (P0 or F1) and three independently propagated F20 lines of wild-type, *mlh-1, pms-2*, and *pole-4* single mutants. **(C)** Number and types of base substitutions observed in the parental (P0) line and 2-3 independently propagated F10 lines wild-type, *pms-2 and pole-4* single, and *pole-4; pms-2* double mutants. **(D)** Number and type of indels observed in the parental (P0) and 2-3 independently propagated F10 lines of wild-type, *pms-2 and pole-4* single, and *pole-4; pms-2* double mutants. **(E)** Average number of base substitutions identified across all individual lines per genotype in their 5’ and 3’ base sequence context in *mlh-1* and *pms-2* single and in *pms-2* single and *pole-4*; *pms-2* double mutants. Error bars represent the standard error of the mean. **(F)** Examples of indel sequence contexts. Sequence reads aligned to the reference genome WBcel235.74 visualized in Integrative Genomics Viewer (Robinson et al. 2011). A 1 bp (left panel) and a 2 bp deletion (right panel) are shown. A subset of sequence reads, which end close to an indel, erroneously aligned across the indel resulting either in wild-type bases (arrow) or base changes (arrowheads). Such wrongly called base substitutions were removed during filtering (Material and Methods) using the deepSNV package (Gerstung et al. 2012; Gerstung et al. 2014).

### Mutation rates and patterns in *pole-4; pms-2* double mutants

To further investigate the role of *pole-4* and the genetic interaction with MMR deficiency, we generated *pole-4; pms-2* double mutants. *pms-2* mutants carried an average of 145 base substitution and 527 indels over 10 generations, roughly half the number we observed in the F20 generation (Fig. 1C and 1D, Supplemental Table S2). In comparison, the number of single base substitutions and indels was increased ~4.4 fold and ~1.4 fold in *pole-4; pms-2* double mutants to an average of 637 and 723, respectively (Supplemental Table S2, Fig. 1C,D). We did not identify any structural variants (SVs) in the genotypes analyzed except for *pole-4*, where a single SV was observed in three F10 mutation accumulation lines (Supplemental Table S2). We could not readily propagate *pole-4; pms-2* beyond the F10 generation, suggesting that a mutation burden higher than ~500–700 single base substitutions (Fig. 1C) in conjunction with 700-750 indels (Fig. 1D) might be incompatible with organismal reproduction. The increased mutation burden of *pole-4; pms-2* double mutants compared to that of *pms-2* and to the wild-type mutation rate of *pole-4* suggests that replication errors occur at increased frequency in the absence of *pole-4* but are effectively repaired by MMR.

Assuming that it takes 15 cell divisions to go through the *C. elegans* life cycle and considering that heterozygous mutations can be lost during self-fertilization (Material and Methods), we calculated a mutation rate per base pair and germ cell division of 1.0 x 10^−9^ (95% CI: 7.96 x 10^−10^ to 1.25 x 10^−9^) for wild-type and 1.19 x 10^−9^ (95% CI: 9.58 x 10^−10^ to 1.45 x 10^−9^) for *pole-4* mutants. In contrast, mutation rates for *mlh-1* and *pms-2* were 7.10 x 10^−8^ (95% CI: 6.86 x 10^−8^ to 7.33 x 10^−8^) and 7.28 x 10^−8^ (95% CI: 7.10 x 10^−8^ to 7.48 x 10^−8^) respectively. *pole-4; pms-2* double mutants exhibited a mutation rate of 1.51 x 10^−7^ (95% CI: 1.45 x 10^−7^ to 1.56 x 10^−7^).

The genome-wide mutation rates observed in the absence of *C. elegans* MutLα proteins MLH-1 and PMS-2 are in line with mutation rates previously determined for *C. elegans* MutS and *S. cerevisiae* MMR mutants (Strand et al. 1993; Yang et al. 1999; Degtyareva et al. 2002; Tijsterman et al. 2002; Denver et al. 2005). However, unlike in mammalian cells (Yao et al. 1999; Baross-Francis et al. 2001), *C. elegans mlh-1* and *pms-2* mutants exhibited almost identical mutation rates and profiles, suggesting that the inactivation of the MutLα heterodimer is sufficient to induce a fully penetrant MMR phenotype consistent with the absence of PMS1 MutLβ and MLH3 MutLγ homologs in *C. elegans*. Our finding that *pole-4* mutants do not show increased mutation rates is surprising given that the deletion of the budding yeast POLE-4 homolog Dpb3 leads to mutation rates comparable to the proofreading-deficient *pol2-4* allele of the Polε catalytic subunit (Aksenova et al. 2010; Lujan et al. 2012). Increased mutation rates have also been reported for proofreading mutants of the Polε catalytic subunit in mice and human and in humans such mutations are associated with an increased predisposition to colorectal cancer (Albertson et al. 2009; Lujan et al. 2012; Palles et al. 2013).

### Distribution and sequence context of base substitutions

We next wished to determine the mutational patterns associated with DNA mismatch repair defects alone and combined with *pole-4* deficiency. T>C and C>T transitions were present more frequently than T>A, T>G, C>A and C>G transversions in *mlh-1* and *pms-2* single and *pole-4; pms-2* double mutants (Fig. 1A,C, Supplemental Table S2). A similar preponderance of T>C and C>T transitions was previously observed in *S. cerevisiae msh2* mutants and in MMR defective human cancer lines (Alexandrov et al. 2013a; Lujan et al. 2014; Supek and Lehner 2015). Analyzing all base substitutions within their 5’ and 3’ sequence context, we found no prominent enrichment of distinct 5’ and 3’ bases associated with T>C transitions in *mlh-1* and *pms-2* single mutants. In contrast, T>A transversions occurred with increased frequency in an A<u>T</u>T context, C>T transitions in a G<u>C</u>N context and C>A transversions in a N<u>C</u>T context (Fig. 1E, Fig. 4A).

Interestingly, > 90% of T>A transversions in an A<u>T</u>T context occurred in homopolymer runs; the majority (> 75%) in the context of two adjoining A and T homopolymers (Supplemental Fig. S1A). An increased frequency of base substitution at the junction of adjacent repeats has also been reported in *S. cerevisiae* MMR mutants, giving rise to the speculation that such base substitutions may be generated by double slippage events (Lang et al. 2013). Moreover, we observed several examples in which one or several base substitutions had occurred that converted a repeat sequence such that it became identical to flanking repeats consistent with polymerase slippage across an entire repeat (Supplemental Fig. S1B-D). Such mechanisms could lead to the equalization of microsatellite repeats, a phenomenon referred to as microsatellite purification (Harr et al. 2000).

While we could not define mutational patterns specifically associated with *pole-4* loss due to the low number of mutations, the profile of *pole-4; pms-2* double mutants differed from MMR single mutants. Most strikingly, in addition to C>T transitions in a G<u>C</u>N context, T>C transitions were generated with higher frequency accounting for >50% of all base changes (Fig. 1C). Among these, T>C substitutions in the context of a flanking 5’ cytosine were underrepresented (Fig. 1E). Interestingly, a higher proportion of T>C changes, not embedded in a defined sequence context, has been reported for MMR-deficient tumor samples containing mutations in the lagging strand polymerase Polδ (Shlien et al. 2015), but not in *S. cerevisiae* and human tumors with a combined MMR and Polε deficiency (Lujan et al. 2014; Shlien et al. 2015). No obvious chromosomal clustering of base substitutions was observed in *pms-2* and *pole-4; pms-2* grown for 10 generations (Supplemental Fig. S2A).

### Sequence context of insertions and deletions associated with MMR deficiency

The majority of mutations observed in *mlh-1* and *pms-2* single and *pole-4; pms-2* double mutants were small insertions/deletions (indels) (Fig. 1B,D). Of these around 90% constituted 1 bp insertions or deletions (Fig. 1B,D, Fig. 2) with most 1bp indels occurring in homopolymer runs (Fig. 1F, Fig. 2B). 2 bp indels accounted on average for 5.5-8.6% of indels observed (Fig. 1B, D, Supplemental Table S2) and affected dinucleotide repeat sequences (Fig. 1F) and homopolymer runs at similar frequency, as recently also reported for MMR mutants in *S. cerevisiae* (Lujan et al. 2015). Trinucleotide repeat instability is associated with a number of neurodegenerative disorders, such as fragile X syndrome, Huntington’s disease and Spinocerebellar Ataxias (Brouwer et al. 2009). Based on our analysis, trinucleotide repeat sequences are present in the *C. elegans* genome at a > 400 fold lower frequency than homopolymer runs (Supplemental Material). We observed between 3 to 7 trinucleotide indels per 10 generations in *mlh-1* and *pms-2* mutants (Supplemental Table S2). However, these occurred predominantly in homopolymer sequences precluding an estimation of mutation rates for trinucleotide repeats.

**Figure 2.**
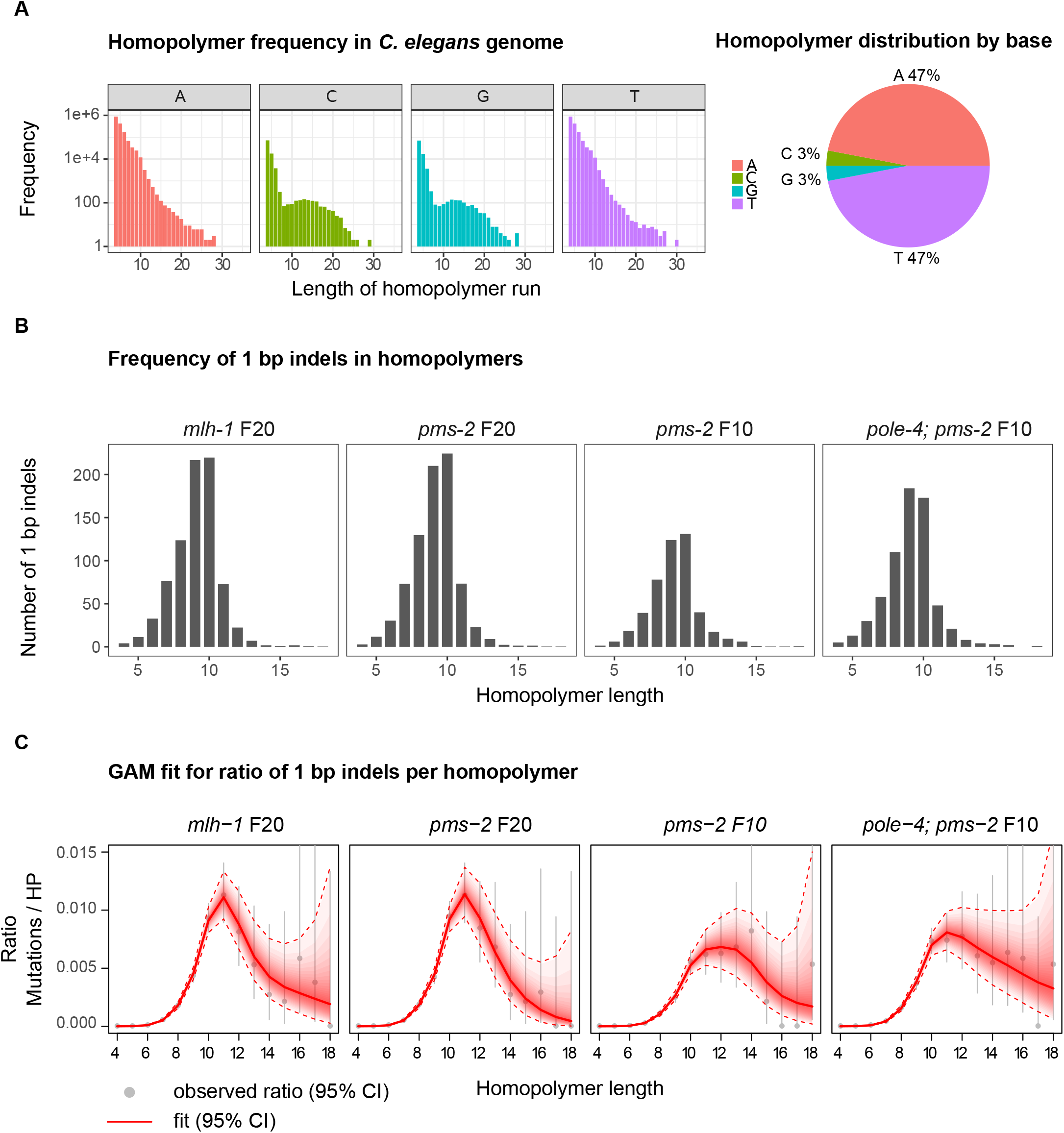
Correlation between homopolymer length and the frequency of +1/−1 bp indels. **(A)** Distribution of homopolymer repeats encoded in the *C. elegans* genome by length and DNA base shown in log_10_ scale (left panel) and the relative percentage of A, C, G and T homopolymers in the genome (right panel). **(B)** Average number of 1 bp indels in homopolymer runs for *mlh-1* F20, *pms-2* F20, *pms-2* F10 and *pole-4; pms-2* F10 mutant lines by homopolymer length. **(C)** Generalized additive spline model (GAM) fit for the ratio of 1bp indels normalized to the frequency of homopolymers (HPs) in the genome. The average frequency observed across three lines is depicted as a grey dot; grey bars indicate the 95% confidence interval. The red line indicates best-fit. Red dotted lines represent the corresponding 95% confidence interval.

Clustering of 1 bp indels was not evident for *pms-2* and *pole-4; pms-2* F10 lines beyond a somewhat reduced occurrence in the center of *C. elegans* autosomes which correlates with reduced homopolymer frequency in these regions (Supplemental Fig. S2B).

### Dependency of 1 bp indel frequency on homopolymer length

Given the high number of indels present in homopolymer repeats we aimed to investigate the correlation between indel frequency and homopolymer length. Overall, we identified 3,433,785 homopolymers in the *C. elegans* genome, ranging in length between 4-35 nucleotides (Fig. 2A, Material and Methods). 47% of all homopolymers each were poly-A or poly-T repeats, their frequencies decreasing with increasing length (Fig. 2A). C and G each comprised 3% of all homopolymers with frequencies decreasing up to a length of 8 bp, plateauing between 8 and 17 bp, and further decreasing for longer homopolymer tracks (Fig. 2A). These findings are consistent with a previous report on >7 bp long homopolymers (Denver et al. 2004). In *C. elegans* MMR mutant backgrounds the frequency of 1 bp indels increased with homopolymer length of up to 9-10 base pairs and trailed off in longer homopolymers (Fig. 2B). Given that the frequency of homopolymer tracts decreases with length (Fig. 2A) we normalized for homopolymer number. To assess the variability of the frequency estimation, we applied an additive model (Material and Methods), which supported a rapid increase in indel frequency in homopolymers up to a repeat length of 10 bases followed by a drop or plateau in indel frequency for longer homopolymers with decreasing confidence (Fig. 2C). Firm conclusions about indel frequencies in homopolymers >13 bp are precluded by the lack of statistical power due to the low numbers of both long homopolymers in the genome and associated indels observed (Fig. 2B). In summary, our data suggest that replicative polymerase slippage occurs more frequently with increasing homopolymer length, with a peak for homopolymers of 10-11 nucleotides, followed by reduced slippage frequency in longer homopolymers. These results are consistent with observations in budding yeast (Lang et al. 2013) and a recent study using human MLH-1^KO^ organoids (Drost et al. 2017).

### Comparison of *C. elegans* MMR patterns to MMR signatures derived from human colorectal and gastric adenocarcinoma samples

To assess how our findings relate to mutation profiles occurring in human cancer we analyzed whole exome sequencing data from the TCGA colorectal adenocarcinoma project (US-COAD) (http://icgc.org/icgc/cgp/73/509/1134) (Cancer Genome Atlas 2012) and the TCGA gastric adenocarcinoma project (US-STAD) (https://icgc.org/icgc/cgp/69/509/70268) (Cancer Genome Atlas Research 2014). These cancer types are commonly associated with MMR deficiency. These datasets contain single nucleotide (SNV) and indel variant calls from 215 and 289 donors, respectively. Having observed high 1bp indel frequencies associated with homopolymer repeats in *C. elegans pms-2* and *mlh-1* mutants (Fig. 1B,D, Fig. 2B), we also considered indels in our analysis of human mutational signatures.

A *t*-distributed stochastic neighbor embedding (t-SNE) representation of the cosine similarities of mutation spectra revealed a distinctive grouping of cancer samples (Fig. 3A; Material and Methods). *De novo* signature extraction across both tumor cohorts combined revealed eight main mutational signatures (Fig. 3B, Material and Methods). Comparing these to existing COSMIC signatures by calculating the similarity score between their base substitution profiles showed that many had a counterpart in the COSMIC database with high similarity (Table 1, Supplemental Fig. S3B), validating our results. We labeled three of the *de novo* signatures as “MMR-1-3”. MMR-1 shares similarity with COSMIC signature 20, MMR-2 with COSMIC signature 15 and MMR-3 with COSMIC signatures 21 and 26 (Table 1, Supplemental Fig. S3B).

**Figure 3.**
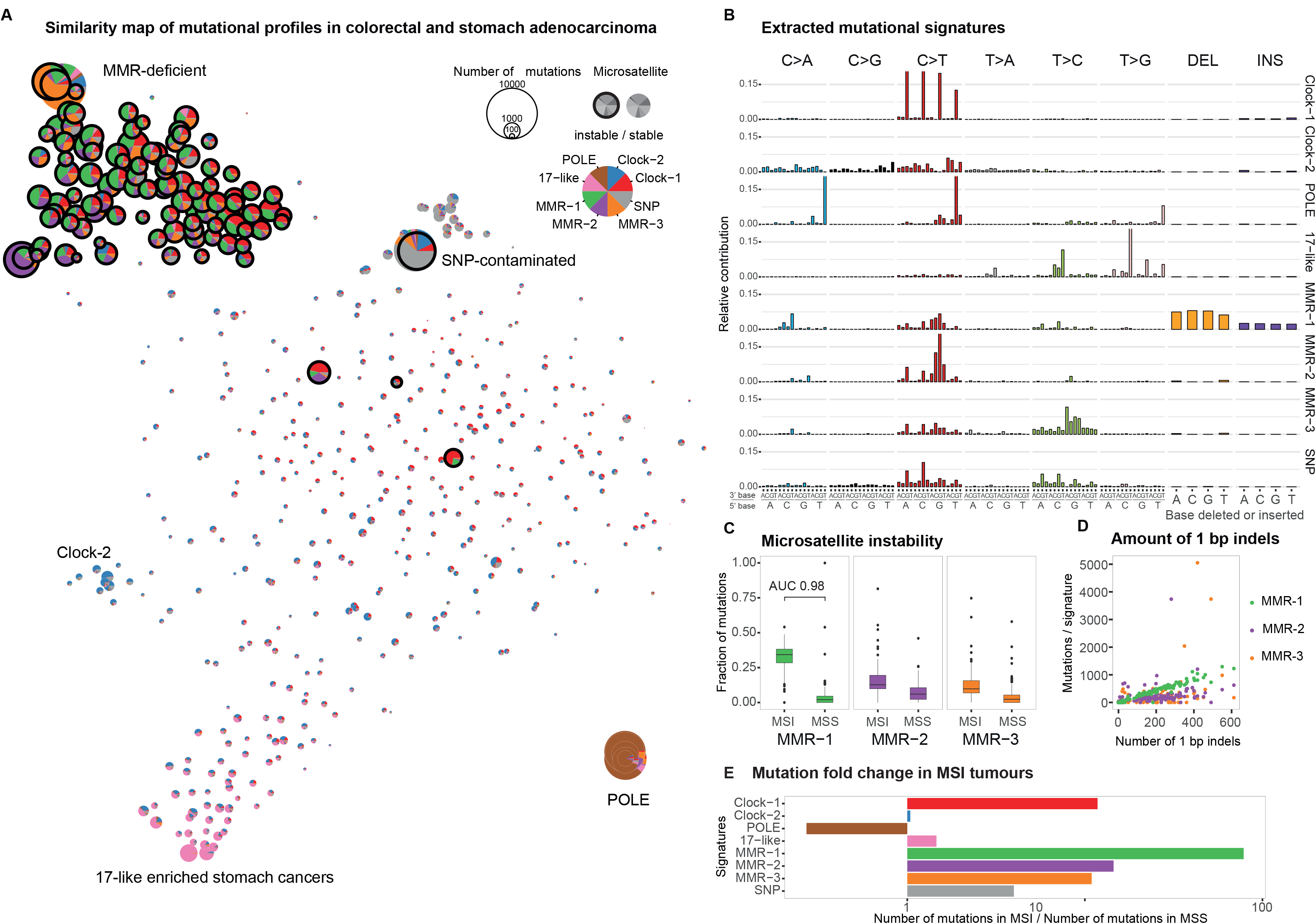
Identification of *de novo* signatures from human colorectal and gastric adenocarcinoma samples (COAD-US and STAD-US projects) and their contribution to samples clinically classified as microsatellite instable (MSI) or microsatellite stable (MSS). **(A)** Two-dimensional representation of the mutational profile composition across cancer samples. The size of each circle reflects the mutation burden. MSI samples are highlighted by a bold, black outline. The color of segments reflects the signature composition. **(B)** Mutational signatures including base substitutions and 1 bp indels derived from the combined COAD-US and STAD-US datasets. **(C)** Relative contribution of MMR-1, MMR-2 and MMR-3 signatures to cancer samples clinically classified as microsatellite instable (MSI) or microsatellite stable (MSS). Box plot with outliers shown as individual filled circles. Area under the curve (AUC) value for MMR-1 contribution indicates the probability of a random MSI sample having higher MMR-1 contribution than a random MSS sample. **(D)** Number of mutations assigned to signatures MMR-1 (green), MMR-2 (purple) and MMR-3 (orange) plotted against the number of 1 bp indels in the same sample. **(E)** Fold change in the average number of mutations assigned to different signatures in MSI samples compared to MSS samples. As expected, the number of POLE related mutations is higher in MSS samples as all POLE-deficient tumours are MSS. Apart from MMR signatures, Clock-1 signature also contributes over 10 times more mutations to MSI samples than to MSS. SNP associated mutations are likely due to unfiltered SNPs that are prevalent in the human population (Supplemental Material).

**Table 1.**
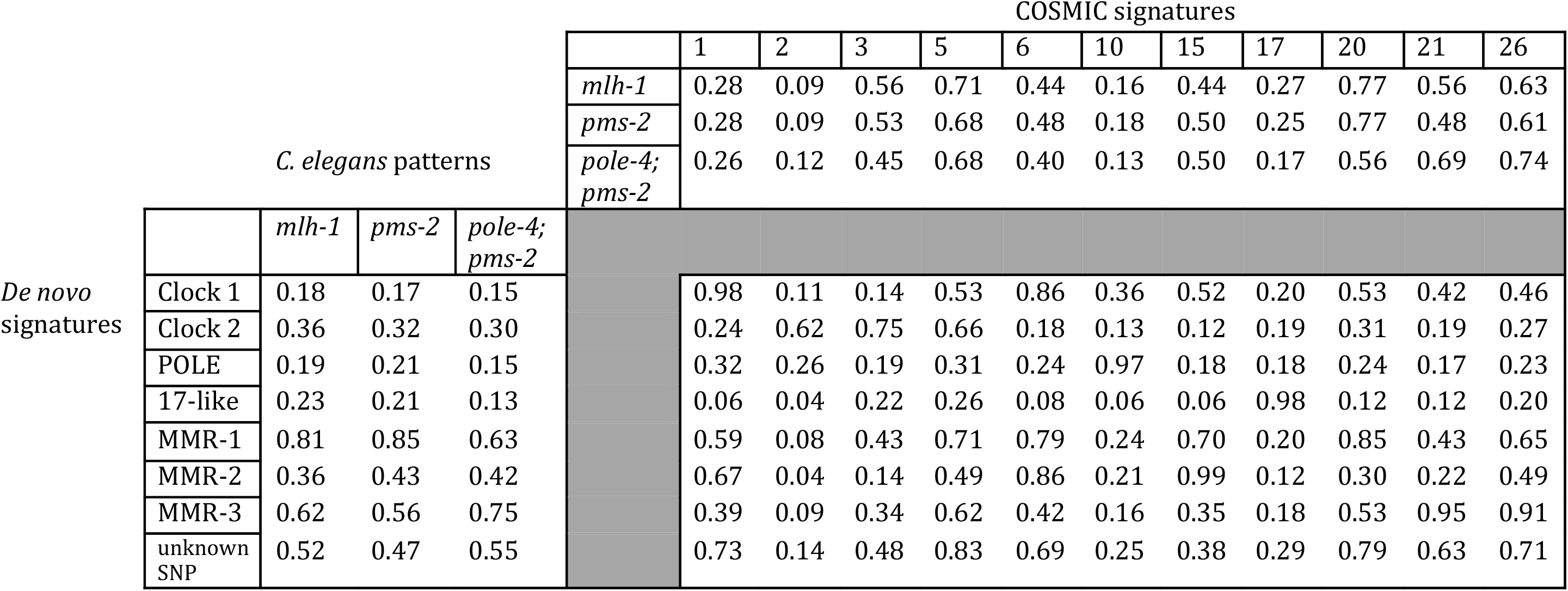
Cosine similarity values for the comparison between humanized *C. elegans* derived MMR mutation patterns with *de novo* signatures and selected COSMIC signatures (both adjusted to human whole-exome trinucleotide frequencies).

Another signature identified in the tumor samples showed characteristics of *POLE* mutations (“POLE”) (Fig. 3B, Supplemental Fig. S3B) (Alexandrov et al. 2013a), http://cancer.sanger.ac.uk/cosmic/signatures). C>T mutations in a CpG base context result from 5meC deamination and are referred to as “Clock-1” (Fig. 3B, Supplemental Fig. S3B). “Clock-2” is present in the majority of samples and likely reflects background mutation rates (Fig. 3A,B). Signature “17-like” is of unknown etiology, predominantly found in stomach cancers, and related to COSMIC signature 17 (Fig. 3B, Supplemental Fig. S3B). Finally, we identified a signature indicative of SNP contamination characterized by a high overlap of somatic mutations with SNPs present in the human population and a lower non-synonymous to synonymous ratio, as expected for germline variants (Fig. 3B, Supplemental Material).

To confirm the link between signatures MMR-1-3 and mismatch repair deficiency, we correlated their occurrence with the microsatellite stability status of samples according to the Stanford TCGA Clinical Explorer (http://genomeportal.stanford.edu/pan-tcga). We considered the samples with MSI-H (microsatellite instable-high) status as MMR-deficient and refer to them as MSI (microsatellite instable), whereas MSI-L (microsatellite instable-low) and MSS (microsatellite stable) samples are considered MMR-proficient and referred to as MSS. 40 out of the 215 (19%) samples in the COAD cohort and 63 out of the 289 (22%) samples in the STAD were assigned MSI status. Comparing MSI/MMR-deficient and MSS/MMR-proficient samples, we found that the fraction of MMR-1, MMR-2 and MMR-3 signatures was strongly enriched in a cluster of microsatellite instable cancer samples (Wilcoxon rank-sum test P-values 1.7 x 10^−51^, 2.7 x 10^−22^ and 9.8 x 10^−29^, respectively) (Fig. 3A top left,C). Consistent with the results from *C. elegans*, 82% of indels in the human cancer samples were 1bp insertions or deletions (25,093 and 30,561 in COAD and STAD cohort, respectively). The majority of these occurred in homopolymer runs at frequencies ranging between 69-72% in COAD and 91-93% in STAD samples (Supplemental Fig. S4B). Of all signatures identified, only MMR-1 highly correlated with the amount of 1 bp indels (Pearson correlation coefficient of 0.99), showing a stable trend compared to other MMR signatures (Fig. 3B,D). MMR-1 signature contribution was a remarkably accurate indicator of MSI/MMR deficiency (AUC of 0.98, 95% confidence interval (CI) 0.97-1).

Individual signatures often represent the most extremes of the mutational spectrum; a typical tumor, however, is usually represented by a linear combination of multiple processes. In the cluster of MSI-H tumors (Fig. 3A, top left) MMR-1 occurs with high relative contribution in the majority of MSI-H tumors, whereas MMR-2 and MMR-3 contributions are generally smaller and more variable across these tumors (Fig. 3A, Supplemental Fig. S5A,B). Interestingly however, the most severely hypermutated samples are largely described by a single signature (Fig. 3A, MMR-2 in purple, MMR-3 in orange). Minor contributions of MMR-2 and MMR-3 in other tumors may be due to the tendency of the signature calculation method to extract signatures predominantly from the most extreme cases, and imparting them on other samples. We considered that MMR-2 and MMR-3 might reflect different substrate specificities of MMR complexes arising from an inactivation of unique subunits of MutSα, MutSβ, MutLα, or MutLγ heterodimers. Investigating MMR gene mutations and methylation status in MSI tumor samples, we only observed few cases of MSH6 (MutSα), MSH3 (MutSβ), PMS2 (MutLα), PMS1 (MutLβ) and MLH3 (MutLγ) high-impact mutations often in combination with inactivation of other MMR genes (Supplemental Material, Supplemental Table S3). Interestingly, we observed an increased number of mutations assigned to the Clock-1 signature in human MMR-deficient samples (Fig. 3A,E). The Clock-1/COSMIC signature 1 is thought to reflect spontaneous 5meC deamination and its conversion to thymine, a mutational process that is thought to be active in all tissues and which correlates with the age at the time of cancer diagnosis (Alexandrov et al. 2013a). Our data suggest a role of MMR in the repair of 5meC deamination-induced mismatches (Bellacosa 2001; Tricarico et al. 2015; Grin and Ishchenko 2016). Notably the most frequent MMR signature, Signature 6, shows high rates of C>T mutations in an NCG context possibly reflecting imperfect delineation of the underlying mutational processes.

A second sample cluster is represented by six tumors (Fig. 3A, bottom right) with most of their mutations falling within the POLE signature (brown). Consistently, these samples also carry pathogenic *POLE* mutations (Supplemental Table S4). Another cluster is formed by a subset of stomach cancer samples carrying a 17-like signature (Fig. 3A, bottom left). Four tumor samples outside of these clusters and dispersed over the similarity map are MSI. These tumors may have acquired MMR deficiency very late in their development.

To compare human and *C. elegans* MMR footprints we first determined mutational patterns from *mlh-1* and *pms-2* single mutants as well as from the *pole-4; pms-2* double mutant (Material and Methods). Mutational patterns of *mlh-1* and *pms-2* mutants were nearly identical with a cosine similarity of 0.97 (Fig. 4A top panels). In contrast the *pole-4; pms-2* mutational pattern showed a different relative contribution of C>T and T>C mutations (Fig. 4A top panels) and displayed a cosine similarity to *mlh-1* and *pms-2* below 0.71 (Supplemental Fig. S6C). We next adjusted for the difference in trinucleotide frequencies in the *C. elegans* genome and the human exome (Fig. 4A bottom panels, Fig. 4B). Comparison of *C. elegans* MMR patterns with known cancer signatures showed the highest similarity of 0.77 to COSMIC signature 20 (Table 1, Supplemental Fig. S7). Of the three human MMR-associated *de novo* signatures, only MMR-1 displayed similarity to *C. elegans* MMR substitution patterns with a cosine similarity of 0.84 to *pms-2* and of 0.81 to *mlh-1* (Table 1, Fig. 4C). A notable difference in the *C. elegans pms-2* and *mlh-1* patterns compared to the MMR-1 signature are a reduced level of C>T mutations in NCG contexts (Fig. 4C, stars) and a high frequency of T>A mutation in an ATT context. The first is likely due to the lack of spontaneous deamination of 5methyl-C, a base modification that is absent in *C. elegans* (Greer et al. 2015), the latter likely due to a higher relative frequency of poly-A and poly-T homopolymers in the *C. elegans* genome versus the human exome (Fig. 2A, Supplemental Fig. S4A). Excluding C>T’s in NCG contexts from the analysis increases the cosine similarity of MMR-1 up to 0.92 for *pms-2* and up to 0.90 for *mlh-1* patterns and reduces the uncertainty in similarity scores (Supplemental Material, Supplemental Fig. S6D). The similarity between *C. elegans* MMR and the human MMR-1 mutational footprints is further supported by the concurrent presence of a large number of 1bp indels. None of the human signatures showed notable similarity to the *pole-4; pms-2* mutation pattern.

**Figure 4.**
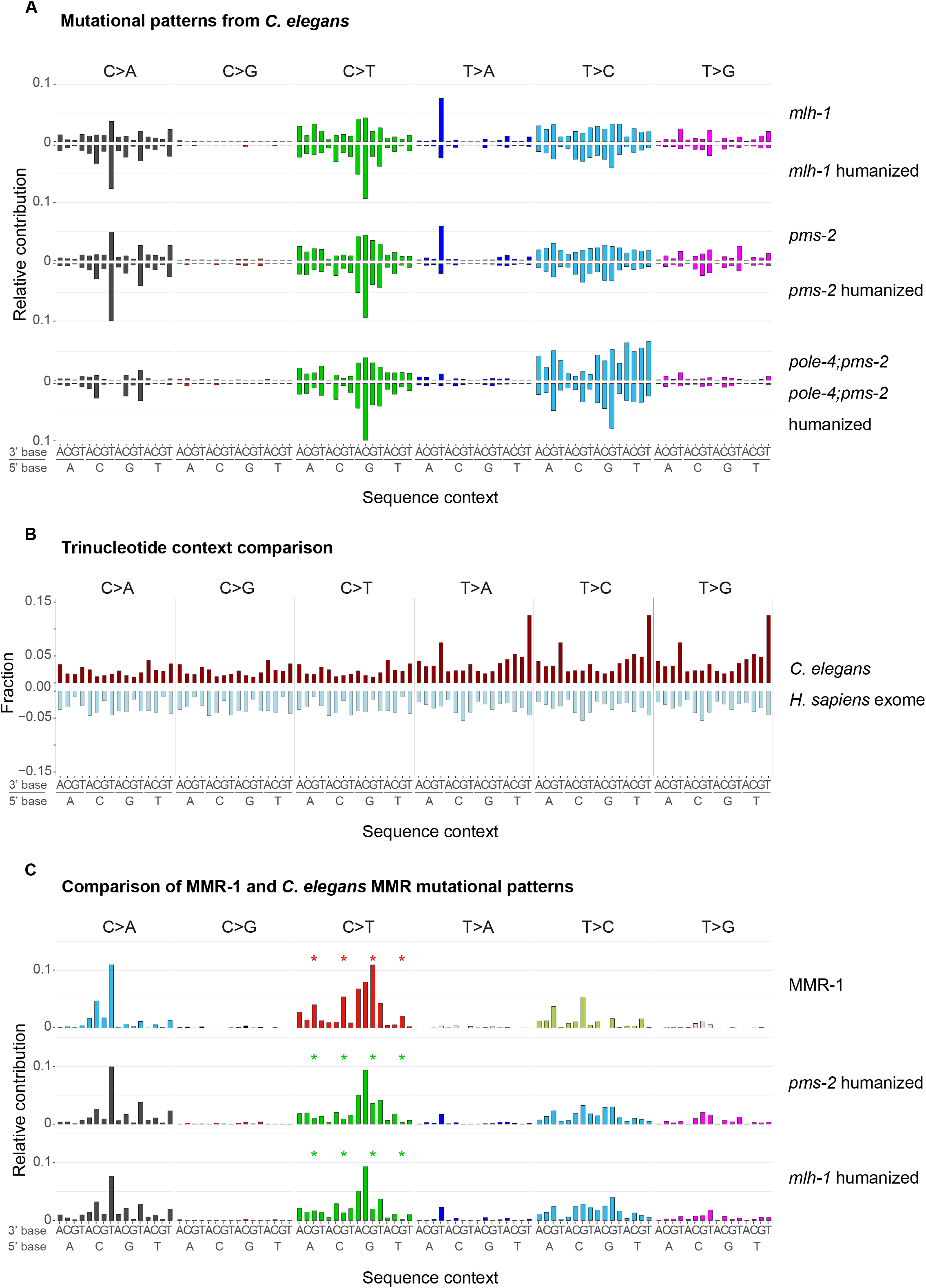
Mutational patterns derived from *C. elegans* MMR mutants and their comparison to the *de novo* human signature MMR-1. **(A)** Base substitution patterns of *C. elegans mlh-1, pms-2* and *pole-4; pms-2* mutants and their corresponding humanized versions (mirrored). **(B)** Relative abundance of trinucleotides in the *C. elegans* genome (red) and the human exome (light blue). **(C)** MMR-1 base substitution signature compared to *pms-2* and *mlh-1* mutational patterns adjusted to human whole exome trinucleotide frequency. Stars indicate the difference in C>T transitions at CpG sites, which occur at lower frequency in *C. elegans*.

## DISCUSSION

MMR-deficient tumors have among the highest mutation rates across cancer types. In line with this observation, we observed an ~70 fold increase in the number of base substitutions in *C. elegans mlh-1* and *pms-2* mutants. This mutation rate is only surpassed by that of the *pole-4; pms-2* double mutant in which mutation rates are further increased 2-3 fold. Genome maintenance is highly efficient as evidenced by a wild-type *C. elegans* mutation rate in the order of 8 x 10^−10^ per base and cell division. It thus appears that DNA repair pathways act highly redundantly, and that it may require the combined deficiency of multiple DNA repair pathways to trigger excessive mutagenesis. Equally a latent defect in DNA replication integrity might only become apparent in conjunction with a DNA repair deficiency. Indeed the increased mutation burden detected in the *pole-4; pms-2* double mutant while no increased mutation rate is observed in *pole-4* alone uncovers a latent role of *pole-4*. It appears that replication errors occur at increased frequency in the absence of *C. elegans pole-4* but are effectively repaired by MMR.

Out of the signatures associated with MMR deficiency in cancer cells, only MMR-1 is related to the mutational pattern found in *C. elegans mlh-1* and *pms-2* mutants. Considering the controlled nature of the *C. elegans* experiment we postulate that MMR-1 reflects a conserved mutational process of DNA replication repaired by MMR. Consistent with this we find that MMR-1 activity is closely linked to MSI status, an established indicator for mismatch repair deficiency. In cases of hypermutation we suggest that akin to the *pole-4; pms-2* double mutant, mutational footprints can be attributed to the failed repair of lesions originating from mutations in DNA repair or DNA replication genes. For instance in MMR defective lines also carrying *POLE* catalytic subunit mutations the mutational landscape is overwhelmed by the POLE signature (Shlien et al. 2015). Likewise it appears possible that the MMR-2 and MMR-3 signatures could be attributed to other mutational processes, which are repaired by MMR and lead to hypermutation under MMR deficiency. Overall, MMR-1 seemingly reflects a ‘basal’ mutational process in both humans and *C. elegans*. In addition, human MMR deficiency also includes an element of failing to repair lesions arising from CpG deamination and leading to C>T mutations, a process absent in *C. elegans* due to the lack of cytosine methylation. The associated human signature, “Clock-1”, together with MMR-1 explains the majority of mutations occurring in MMR defective cancers not apparently affected by hypermutation.

Matching mutational signatures to DNA repair deficiency has a tremendous potential to stratify cancer therapy tailored to DNA repair deficiency. This approach appears advantageous over genotyping marker genes, as mutational signatures provide a read-out for cellular repair deficiency associated with either genetic or epigenetic defects. Following on from our study we expect that analyzing DNA repair defective model organisms and human cell lines, alone or in conjunction with defined genotoxic agents, will contribute to a more precise definition of mutational signatures occurring in cancer genomes and to establishing the etiology of these signatures.

## MATERIAL AND METHODS

### *C. elegans* maintenance and propagation

*C. elegans* mutants *pole-4(tm4613) II*, *pms-2(ok2529) V* and *mlh-1(ok1917) III* were backcrossed 6 times against the wild-type N2 reference strain TG1813 (Meier et al. 2014) *pole-4 II; pms-2 V* double mutants were generated as described in Supplemental Material. Strains were grown for 10 or 20 generations and genomic DNA was prepared as described (Supplemental Material) (Meier et al. 2014).

### DNA sequencing, variant calling and post-processing

Illumina sequencing, variant calling and post-processing filters were performed as described (Meier et al. 2014) with the following adjustments. WBcel235.74.dna.toplevel.fa was used as the reference genome (http://ftp.ensembl.org/pub/release-74/fasta/caenorhabditis_elegans/dna/). Alignments were performed with BWA-mem and mutations were called using CaVEMan, Pindel and BRASSII (Ye et al. 2009; Stephens et al. 2012), each available at (https://github.com/cancerit). Post-processing of mutation calls was performed across a large dataset of 2202 sequenced samples using filter conditions described (Supplemental Material) (Meier et al. 2014). An additional filtering step using deepSNV (Gerstung et al. 2012; Gerstung et al. 2014) was used to correct for wrongly called base substitutions, events due to algorithm-based sequence misalignment of ends of sequence reads covering 1bp indels (see Fig. 2C).

### Estimating mutation rates

Mutation rates were calculated using maximum likelihood methods, assuming 15 cell divisions per generation (Meier et al. 2014), and considering that mutations have a 25% chance to be lost, a 50% chance to be transmitted as heterozygous, and a 25% chance to become homozygous, thus becoming fixed in the line during each round of *C. elegans* self-fertilization. Wild-type, *pole-4* and *pms-2* mutation rates were calculated from mutations observed across F10 and F20 generations.

### Analysis of homopolymer sequences in *C. elegans* and human cancer samples

Homopolymers, di- and tri-nucleotide runs encoded in the *C. elegans* genome, defined here as repetitive DNA regions with a consecutive number of identical bases or repeated sequence of n≥4, were identified from the reference genome WBcel235.74 using an in house script (Supplemental Data Analysis, https://gerstung-lab.github.com/MMR) based on R packages Biostrings and GenomicRanges (Lawrence et al. 2013; Pagès et al. 2016) Overall, 3,433,785 homopolymers, 25,126 dinucleotide and 7,615 trinucleotide repeats were identified. Matching the genomic position of 1 bp indels observed in *pms-2* and *pole-4; pms-2* mutants to the genomic positions of homopolymers, we defined 1bp indels present in homopolymer runs and the length of the homopolymer in which they occurred.

Generalised additive models with a spline term were used to correlate the frequency of 1 bp indels occurring in homopolymer runs with the frequency of the respective homopolymer in the *C. elegans* genome. An average of ~ 0.5 1bp indels arising in homopolymer sequences was observed in 101 wild-type lines of different generations, indicating the frequency with which such events might occur in wild-type or as amplification artefacts during sequencing (Supplemental Material). Consistent with COAD and STAD variant calling, homopolymer frequencies and coordinates in the human exome were calculated based on the GRCh37 human reference genome build and the coordinates of well-covered fraction of whole-exome sequencing regions as reported by Agilent SureSelect V5 Human All Exon (https://earray.chem.agilent.com/earray/). Overall, we identified 976,390 homopolymers in the human exome, which ranged from 4 to 35 basepair in length (Supplemental Fig. S4A). A more recent genome build, GRCh38, does not differ in the composition of coding regions. Therefore the analysis of homopolymer frequencies is valid using both assembly versions.

### *De novo* signature extraction from human cancer samples

Variant calls for whole-exome sequencing data from the colorectal adenocarcinoma (COAD-US) and gastric adenocarcinoma (STAD-US) projects were taken from ICGC database (http://icgc.org). Mutational counts and contexts were inferred from variant tables using MutationalPatterns R package (Blokzijl et al. 2016). After combining indels and substitutions into vectors of length 104, we extracted the signatures using the Brunet NMF with KL-divergence as implemented in (Blokzijl et al. 2016), which is equivalent to a Poisson NMF model. The number of signatures was determined based on the saturation of both the Akaike Information Criterion (AIC) (Akaike 1973) and the residual sum of squares (RSS) (Supplemental Fig. S3A, Supplemental Material). Similarity between signatures was calculated via cosine similarity:

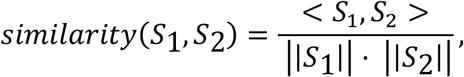

where < *S*_1_, *S*_2_ > is a scalar product of signature vectors. When compared to 96-long substitution signatures, indels were omitted from 104-long *de novo* signatures.

Stochastic nearest neighbor representation (*t*-SNE) (van der Maaten and Hinton 2008) was obtained using R-package “tsne” (Donaldson 2016) using the cosine similarity as distance measure between mutational profiles. In order to confirm the link between signatures MMR-1-3 and MMR deficiency, we defined MMR-deficient samples as those annotated as MSI-H (microsatellite instable high) in TCGA Clinical Explorer (Lee et al. 2015). Relative contributions of every signature to the samples from the combined dataset were tested for association with MSI/MSS status using one-tailed Wilcoxon rank sum test. All p-values were adjusted for multiple testing correction using Bonferroni procedure.

### Comparison of *C. elegans* mismatch repair mutation patterns to cancer signatures

To extract the signatures of individual factors from respective *C. elegans* samples, we used additive Poisson model with multiple factors for every trinucleotide context and indel type and calculated maximum likelihood estimates for every signature (Supplemental Material). For comparison of *C. elegans* and human mutational signatures, signatures acquired in *C. elegans* were adjusted by multiplying the probability for 96 base substitutions by the ratio of respective trinucleotide counts observed in the human exome (GRCh37, counts pre-calculated in (Rosenthal et al. 2016) to those in the *C. elegans* reference genome (WBcel235). Indels were not included in the comparative analysis as they required adjustment for both base and homopolymer content. COSMIC signatures were also adjusted to exome nucleotide counts as they were mostly derived from whole exomes (Alexandrov et al. 2013a; Alexandrov et al. 2013b) and the comparison of *de novo* signatures to COSMIC is more valid in exome space. All signatures were further normalized so that the vector of probabilities sums up to 1 (Supplemental Table S5). For mutational signature comparison a cosine similarity of 0.80 was considered a threshold for “high” similarity (Supplemental Material, Figure S6A,B).

### Data Analysis

R (https://www.R-project.org, R Core Team 2017) scripts used to analyse *C. elegans* and cancer datasets are available in Supplemental Data Analysis and on Github under https://gerstung-lab.github.com/MMR.

## DATA ACCESS

Sequencing data from this study have been submitted to the NCBI Sequence Read Archive (SRA; http://www.ncbi.nlm.nih.gov/sra) under accession number SRP020555.

## ACKNOWLEDGEMENTS

This work was supported by a Wellcome Trust Senior Research award to AG (090944/Z/09/Z) and by a Wellcome Trust Strategic Award (101126/B/13/Z). B.M., N.V.V., P.J.C., M.G., and A.G. are members of the Wellcome Trust funded COMSIG (Causes of Mutational SIGnatures) consortium. P.S. and the Data Analysis Group, Dundee, were funded by the “Wellcome Trust Technology Platform” Strategic Award 097945/Z/11/Z. We thank the Mitani laboratory funded by the National Bio-Resource Project, Japan, and the Caenorhabditis Genetics Center funded by the NIH Office of Research Infrastructure Programs (P40 OD010440) for providing strains. We also thank Steve Hubbard, University of Dundee, for help with statistics.

